# Human CNS-3D organoids predict clinical seizure liability from calcium network activity

**DOI:** 10.64898/2026.05.20.726675

**Authors:** Andrew S. LaCroix, Nicholas S. Coungeris, Victoria Alstat, Corey Rountree, Paolo Botta, Muhammad Maaz, Christopher M. Butt

## Abstract

Drug-induced seizures remain a major safety concern in drug development, yet human seizure liability is difficult to predict using conventional preclinical models. Here, we evaluated whether spontaneous calcium network activity in human induced pluripotent stem cell-derived CNS-3D Brain Organoids could predict clinically observed seizure risk across a pharmacokinetically anchored drug set. CNS-3D organoids contained neuronal and astrocytic populations, expressed neuroactive receptor and ion-channel gene programs that aligned with human cortical tissue, and exhibited reproducible spontaneous calcium oscillations across production batches. A retrospective drug panel of 66 small-molecule drugs was assembled from human clinical evidence, including 30 seizure-associated drugs and 36 comparator drugs without documented clinical seizure liability. Drugs were tested across concentration ranges anchored to reported clinical C_max_, and calcium time-series responses were integrated with chemical structure features using a machine-learning workflow. The final model predicted clinical seizure liability with an AUROC of 0.872, achieving 83.3% sensitivity and 88.9% specificity in drug-level cross-validation. Model scores also stratified seizure-associated drugs by clinical context and prevalence, suggesting that CNS-3D activity profiles capture clinically meaningful differences in seizure risk. Compared with published *in vitro* and preclinical seizure-liability models, CNS-3D organoid-based predictions showed improved balanced sensitivity and specificity. These findings support high-throughput calcium profiling in human CNS-3D organoids as a scalable, exposure-aware platform for predicting human seizure liability and contributing functional human data to neuro-safety assessment.

## Introduction

Seizure liability remains a persistent safety concern in drug discovery and development, contributing to acute, risk-management decisions in the clinic as well as attrition and delayed progression of potential therapeutics^1^. Drug-induced seizures may occur during therapeutic exposure, overdose situations, or during abrupt discontinuation and withdrawal^2^. Thus, any association of a drug with seizure affects dose selection, labeling, and the overall risk–benefit assessment of a candidate drug.

Despite its importance, seizure liability remains difficult to predict before human exposure. Current approaches rely substantially on observations from rodent and non-rodent toxicology studies, brain-slice preparations^3,4^, electrophysiology assays^1,5^, and other reductionist *in vitro* systems^6,7^. Unfortunately, each of these approaches is limited by species differences, assay throughput, endpoint selection, or incomplete representation of network behavior in the human brain^8^.

Neural systems derived from human induced pluripotent stem cells (iPSC) offer a complementary strategy for evaluating neurotoxicity within the contexts of human genetics and human cellular function. Three-dimensional neural models, including brain microphysiological systems, organoids^9-11^, and spheroids^12^, can incorporate multiple neural cell types and support spontaneous network activity that is not readily captured in simpler cellular systems. The CNS-3D Brain Organoids that we have developed from human iPSCs extend this approach by not only containing neuronal and glial populations, but these cell populations also exhibit spontaneous, coordinated calcium oscillations^13^. Importantly, these calcium oscillations are suitable for high-throughput functional screening^14,15^.

Calcium oscillations are particularly useful for neuro-safety assessment because each transient reflects a coordinated burst of neuronal firing and intracellular calcium flux. The disruption of these network bursts has been correlated to neurotoxicity^16,17^. The signals can be recorded in multiwell formats with fluorescence imaging plate readers, enabling quantitative measurement of network-level responses across dose and time^13-17^. Critically, prior CNS-3D work has shown that calcium-activity metrics can detect drug-induced functional disruption at concentrations lower than those that cause decreased viability or morphological degeneration^13^. This dynamic is central to the utility of the assay: functional perturbations can be detected before cellular injury becomes apparent.

Prior studies have established that human iPSC-derived 3D neural cultures respond to pharmacological modulators of excitatory and inhibitory signaling^13-17^. Multiparametric analysis of this signaling can then be used for phenotypic screening^15^ and neurotoxicity profiling^16,17^. Related work has also benchmarked microelectrode array and calcium oscillation assays across rodent, human 2D, and human 3D neuronal systems for de-risking seizure liability^5-7,16,17^. However, much of this literature has emphasized tool drugs, mechanistic pharmacology, or limited reference sets. Fewer studies have paired high-throughput, human 3D functional measurements with pharmacologically relevant exposure ranges and clinical outcome labels. Moreover, most studies have not been performed at large enough scales to support prediction of human seizure liability for a given drug.

The work reported here evaluated whether dose-dependent calcium activity in CNS-3D Brain Organoids could be used to predict clinically observed seizure risk. We assembled a retrospective, clinically annotated dataset of 66 small-molecule drugs, including drugs associated with therapeutic-range seizures, overdose-related seizure events, and CNS-penetrant comparator drugs without documented seizure liability. Drug exposures were anchored to human pharmacokinetic data and tested across concentration ranges that spanned 0.1× to 100× the maximum plasma concentrations (C_max_) observed in clinical samples. The functional calcium time-series responses to each drug were then integrated with molecular structure information using a machine-learning workflow to generate drug-level neurotoxicity scores.

This study links CNS-3D network activity to clinically observed seizure outcomes. By combining a high-throughput human organoid assay with clinically grounded drug annotation and machine-learning analysis, CNS-3D functional profiling provides a scalable approach for identifying seizure liability and supporting neuro-safety decisions within a broader, weight-of-evidence framework.

## Results

### CNS-3D organoids exhibit defined cortical cell populations, stable neuroactive gene expression, and spontaneous calcium oscillations

CNS-3D organoids were first characterized to define the cellular, transcriptomic, and functional properties of the assay system (Fig. 1). Immunocytochemical labeling of neurons (MAP2) and astrocytes (GFAP) shows a relative balance in these two major co-differentiated cell types (Fig. 1a). Furthermore, single-nucleus RNA-seq identified six recurrent cell populations: mature glutamatergic neurons, immature glutamatergic neurons, GABAergic neurons, astrocytes, and two small progenitor cell populations (Fig. 1b).

**Figure 1.**
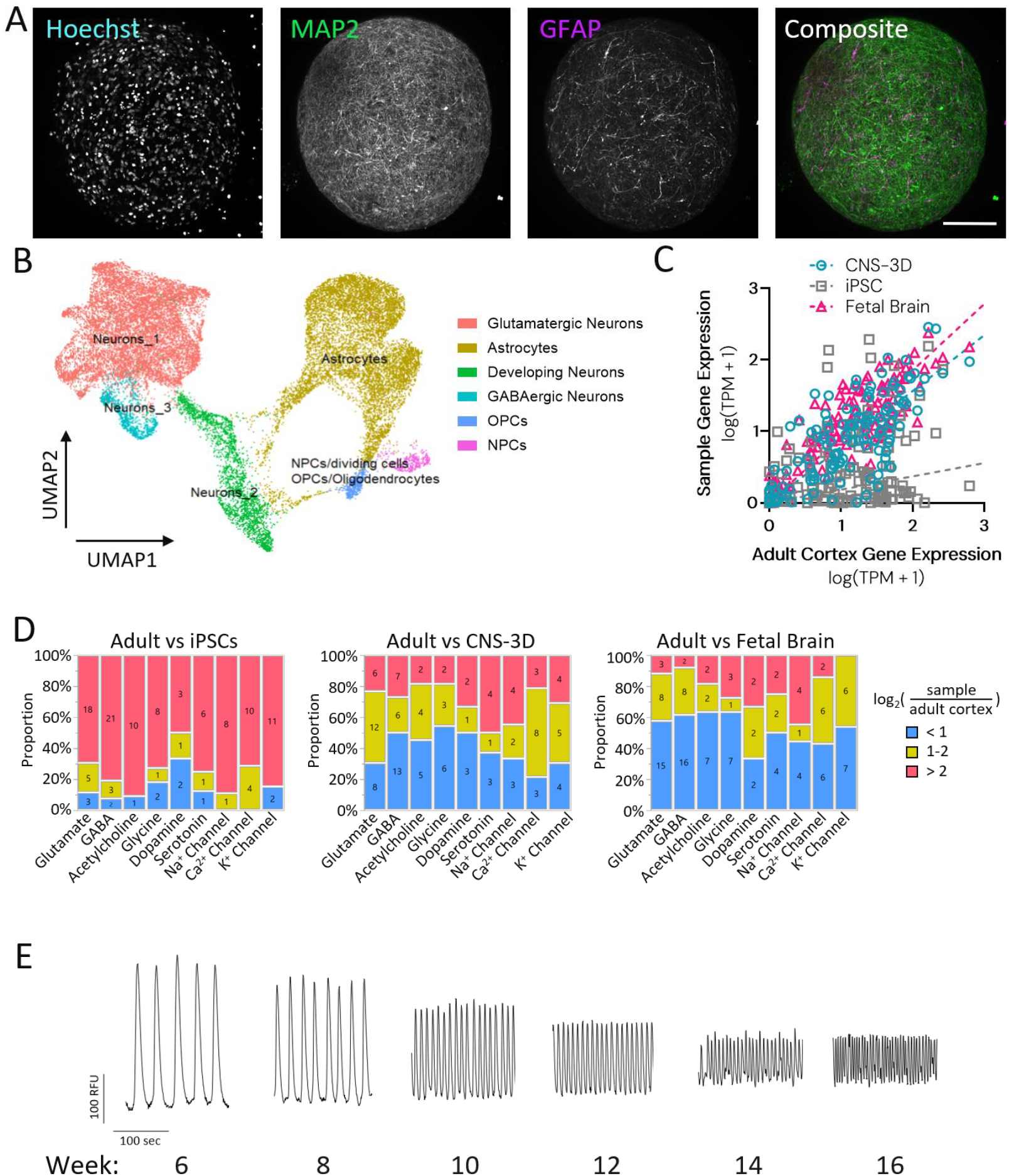
CNS-3D Brain Organoid cell populations, gene expression profiling, and spontaneous calcium measurements. (A) Representative immunofluorescence images of CNS-3D organoids stained for nuclei (Hoechst), neurons (MAP2), and astrocytes (GFAP); scale bar 200 µm. (B) UMAP visualization of single-nucleus RNA-seq profiles from CNS-3D organoids; n=64 organoids pooled per sample from n=3 independent experiments. (C) Correlation of targeted neuroactive gene expression in CNS-3D organoids, iPSCs, and fetal whole brain relative to adult cortex; expression values are shown as log_10_(TPM + 1). (D) Proportion of genes in major neurotransmitter and ion channel families categorized as closely aligned (<1 log_2_ unit), moderately aligned (1–2 log_2_ units), or divergent (>2 log_2_ units) relative to adult cortical expression. (E) Representative FLIPR calcium traces from CNS-3D organoids between 6 and 16 weeks of differentiation.

Gene expression profiles of CNS-3D organoids at 10 weeks of differentiation were compared with those of source iPSCs, human adult cortex, and human fetal brain mRNA samples. Gene-level analysis confirmed expression of neurotoxicity- and seizure-associated signaling pathways in CNS-3D organoids, including glutamatergic and GABAergic receptor families, voltage-gated sodium, potassium, and calcium channels, and other neurotransmitter receptor systems (Supplemental Table 1). CNS-3D organoid expression profile showed an overall strong correlation with adult cortex (Spearman ρ = 0.715), approaching the correlation observed between adult cortex and fetal brain (ρ = 0.803). In contrast, undifferentiated iPSCs showed substantially weaker correlation (ρ = 0.277) with adult cortical gene expression (Fig. 1c). To evaluate the magnitude of expression differences, the log_2_ fold change (log_2_FC) relative to adult cortex was calculated for each sample and gene. Genes were categorized as closely aligned (<1 log_2_ unit), moderately aligned (1–2 log_2_ units), or divergent (>2 log_2_ units) relative to adult cortical expression. Compared with the iPSC source, CNS-3D organoids and fetal brain showed substantially closer alignment with adult cortex, with most genes across major neurotransmitter and ion channel families falling within one or two log_2_ units of adult cortical expression (Fig. 1d).

The spontaneous activity of the system was assessed with FLIPR-based calcium imaging. CNS-3D organoids exhibited spontaneous, coordinated calcium oscillations that evolved over the course of differentiation. Coordinated network bursts first appeared after approximately 6 weeks of differentiation and continued to increase in frequency through at least 16 weeks of maturity (Fig. 1e). Past 12 weeks of differentiation, spontaneous activity reached levels that made height and shape features too difficult to quantify in a rigorous manner. Therefore, assays were restricted to organoids <12 weeks of maturity. Together, these findings establish CNS-3D organoids as a human iPSC-derived cortical system that contains relevant cell populations, neuroactive signaling pathways, and coordinated, spontaneous, functional activity.

### CNS-3D organoid production methods yield stable cellular composition, gene expression, and functional activity across batches

We next assessed the reproducibility of CNS-3D organoid composition and functional activity across independent organoid batches and screening experiments (Fig. 2). The cellular populations identified via single-nucleus RNA-seq (Fig. 1b) were proportionally consistent across three independent organoid batches (Fig. 2a). PCA of selected neurotoxicity-associated genes showed tight clustering of CNS-3D organoid samples between 6 and 10 weeks of differentiation, indicating the biological stability of the organoids once spontaneous activity is observed (Fig. 2b). CNS-3D organoids clustered with fetal brain and adult cortex references along the first principal component, which accounted for nearly two-thirds of the expression variance. Representative activity traces showed coordinated spontaneous oscillations across organoids from the same plate, with similar burst frequency, height, and structure over the 10-minute recording window (Fig. 2c). Finally, plates produced as part of this work were tested for their ability to respond to both excitatory (4-aminopyridine, K+ channel blocker) and inhibitory (muscimol, GABA_A_ receptor agonist) drugs. Across four independent experiments, CNS-3D organoids met prespecified functional QC criteria, with clear separation between active and inhibited network states (Z′ > 0.5, Fig. 2d) and preserved excitatory responsiveness, as shown by robust 4-aminopyridine–evoked increases in spontaneous bursting (>150% normalized peak frequency, Fig. 2e). Together, these data demonstrate that CNS-3D organoids maintain stable cellular composition, gene expression profiles, and spontaneous functional activity across independent experiments.

**Figure 2.**
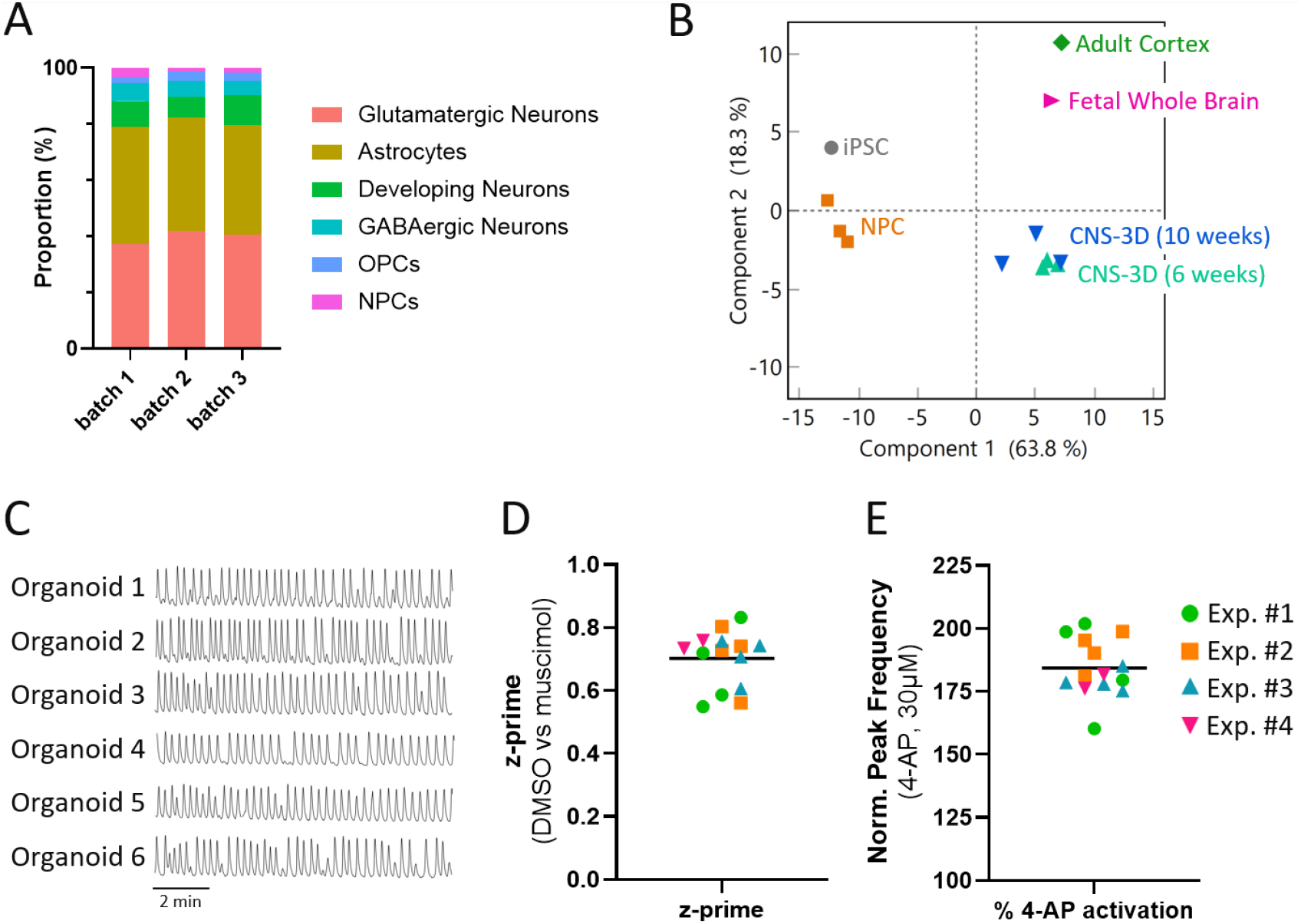
Reproducibility of CNS-3D organoid composition, gene expression, and calcium activity. (A) Proportions of cell populations identified by single-nucleus RNA-seq across CNS-3D organoid batches; n=64 organoids pooled per sample from n=3 independent experiments. (B) PCA of bulk RNA-seq profiles of CNS-3D organoids at 6 and 10 weeks of differentiation compared to iPSC, NPC, adult cortex, and fetal whole-brain reference samples; n=3 independent batches of NPCs and CNS-3D organoids. (C) Representative spontaneous calcium activity traces from six CNS-3D organoids recorded over a 10-minute recording. (D) Z′ values for vehicle versus 10 µM muscimol treated organoids and (E) normalized peak frequency following 30 µM 4-aminopyridine treatment; 100% represents vehicle-treated organoid activity; n=4 independent experiments comprising n≥3 plates each and n≥16 organoids per plate; data points represent individual screening plates; horizontal lines indicate mean values.

### Convulsants produce distinct waveforms of calcium activity in CNS-3D organoids

We next tested whether the spontaneous calcium activity in CNS-3D organoids could capture seizure-relevant network perturbations induced by convulsants (Fig. 3). 4-aminopyridine produced a marked increase in peak frequency, consistent with hyperexcitability following potassium channel blockade (Fig. 3a). Bicuculline, a GABA_A_ receptor antagonist, produced only a moderate increase in peak frequency (Fig. 3b). In contrast, kainic acid, an activator of ionotropic glutamate receptors, increased oscillation frequency while decreasing peak height (Fig. 3c). Together, these diverse phenotypes indicate that clinical seizure liability may be encoded not only in changes in oscillation frequency, but also in the broader structure of the calcium waves.

**Figure 3.**
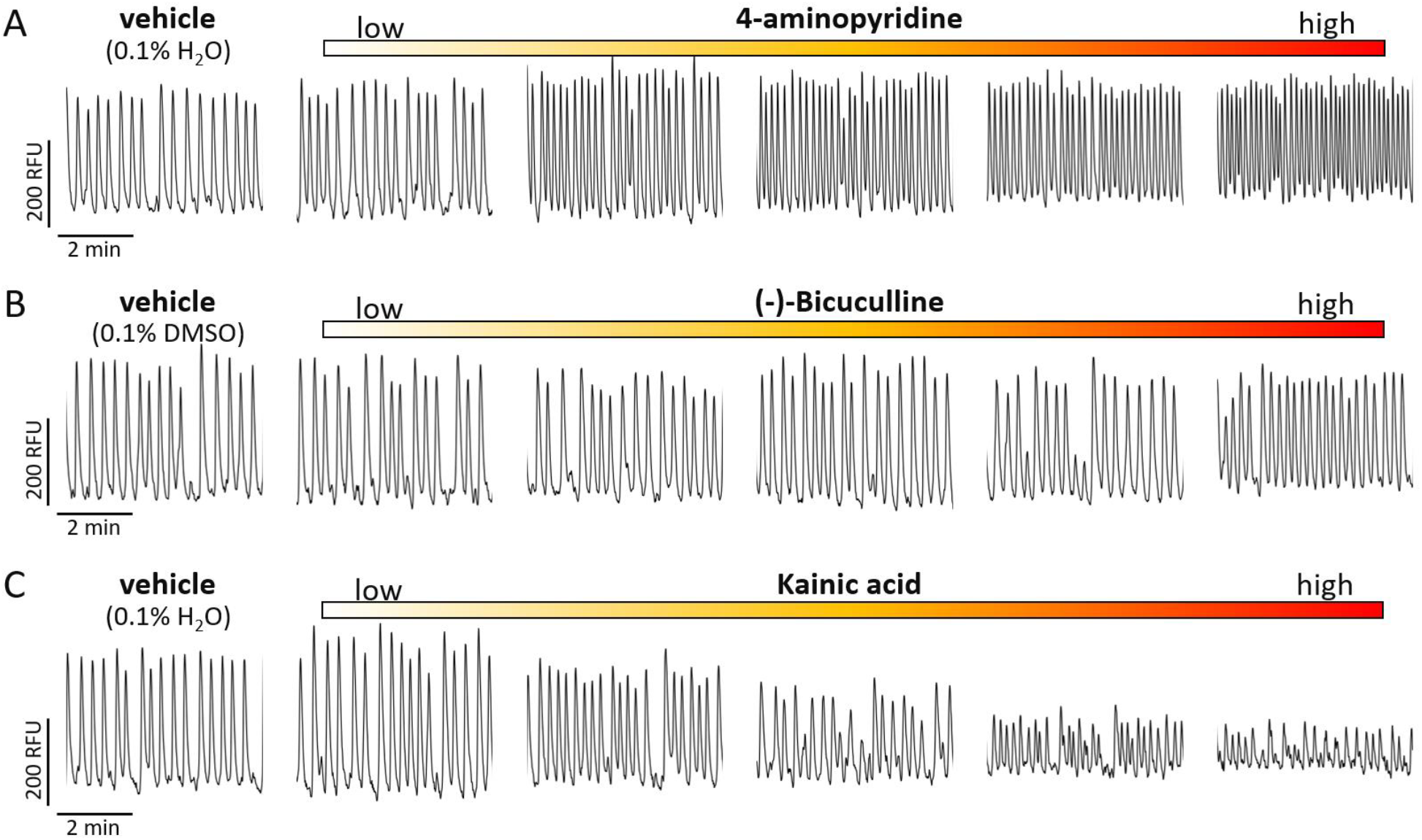
Dose-dependent functional responses to canonical convulsants in CNS-3D organoids. Representative FLIPR calcium activity traces from CNS-3D organoids treated with vehicle and increasing concentrations of canonical convulsant drugs (A) 4-aminopyridine (0.3, 1, 3, 10, and 30 µM), (B) (-)-bicuculline (1, 3, 10, 30, and 100 µM) and (C) kainic acid (0.03, 0.1, 0.3, 1, and 3 µM). Traces show fluorescence intensity over time from representative wells after 2 hours of drug exposure.

### Clinically safe and seizure-associated drugs were selected to reflect therapeutic exposures and enable phenotype stratification

Since convulsants produced multidimensional calcium responses rather than a simple change in frequency, we assembled a drug set anchored on human outcomes in the clinic. The dataset comprised 66 small-molecule drugs evaluated retrospectively against outcomes from 120,551 patients, including 30 seizure-associated drugs and 36 comparator drugs without documented clinical seizure liability^18^. This set was then used to test whether the activity profiles of the organoids contain data that are predictive of human seizure liability (Supplemental Table 2).

Drug inclusion required well-documented clinical outcomes, defined chemical structures represented by canonical SMILES strings, and pharmacokinetic information sufficient for exposure anchoring. Drugs were selected to span diverse mechanisms in both the seizure-associated and comparator classes, including pro-convulsants, glutamate agonists, GABA antagonists, ion channel modulators, paradoxical anticonvulsants, and drugs with other primary mechanisms.

Each drug was tested across a seven-point, half-log, concentration-response series that spanned approximately 0.1x to 100x the reported clinical C_max_. The seizure-associated class included drugs linked to seizures at therapeutic exposure levels as well as drugs associated with overdose-related seizure events. Together, this evaluation provided an exposure-matched dataset for determining whether CNS-3D calcium activity profiles are predictive of human seizure liability.

### Integrated CNS-3D organoid model predicts clinical seizure liability with high sensitivity and specificity

We next developed a machine learning workflow to convert dose-dependent calcium responses into drug-level neurotoxicity scores. Calcium recordings were processed using a feature-extraction workflow that generated functional descriptors from the shape and temporal structure of each recording. Chemical structure was encoded separately based on canonical SMILES strings. Based on mutual information scores, functional time-series features showed the strongest association with seizure classification, whereas SMILES-derived chemical descriptors alone retained weaker but measurable signal. The 120-min post-exposure recording produced the strongest performance among tested time points and was selected for final model training and evaluation.

The final integrated model achieved an AUROC of 0.872 for clinical seizure liability prediction (Fig. 4a). At the threshold selected by Youden’s J statistic, sensitivity was 83.3% and specificity was 88.9%, indicating high, balanced performance in both identifying seizure-associated drugs and confirming the safety profile of known CNS-safe drugs. We examined whether continuous model scores captured a continuum of seizure severity within the seizure-associated class. Drugs associated with overdose-related seizures had significantly higher neurotoxicity scores than drugs associated with therapeutic-range seizure events (Fig. 4b). Higher model scores were also associated with greater prevalence of seizures observed in the clinic (Fig. 4c). These findings demonstrate that CNS-3D functional profiling combined with machine learning predicts clinical seizure liability with high sensitivity and specificity. Furthermore, the model generates continuous scores that could be indicative of differences in seizure mechanism and risk that are clinically meaningful.

**Figure 4.**
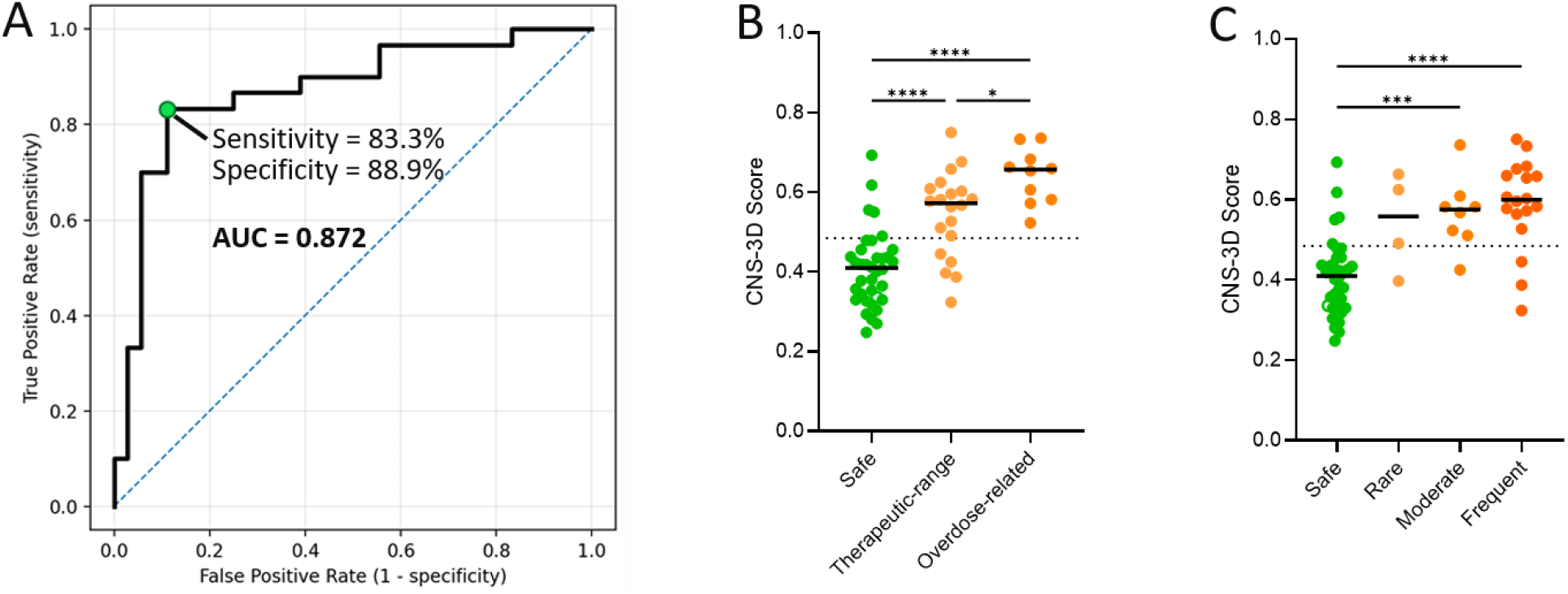
Performance of CNS-3D organoids in predicting clinical seizure liability for small-molecule drug set. (A) ROC curve for seizure liability prediction with AUC of 0.872; optimal threshold is highlighted in green and provides a sensitivity of 83.3% and specificity of 88.9%; blue dotted line indicates the performance expected under random classification. (B) Predicted scores for small-molecule drug panel grouped by seizure type. (C) Predicted scores for small-molecule drug panel grouped by seizure prevalence. Statistical significance assessed using 1-way ANOVA followed by Tukey’s post hoc test; ^*^ p < 0.05, ^**^ p < 0.01, ^***^ p < 0.001, ^****^ p < 0.0001.

### CNS-3D organoid-based predictions outperform existing *in vitro* and preclinical seizure models

We compared the performance of the CNS-3D organoid model with other published methods for predicting seizure liability, including multi-electrode array (MEA) recordings in rat and human 2D systems, calcium oscillation assays in rat, human 2D, and human 3D systems, human ion-channel assays, and prior CNS-3D calcium-based models (Fig. 5)^5-7,16,17^. Across comparator studies, sensitivity and specificity varied substantially. Animal-derived models provided good sensitivity, but poor specificity, indicating a tendency towards false positive classification. Predictions based on human 2D mono- or co-cultures tended to show a tradeoff between identifying seizure-associated drugs and avoiding misclassification of clinically safe drugs. Higher sensitivity models tended to have lower specificity and vice versa. CNS-3D organoid-based predictions broke this trend and achieved best-in-class sensitivity and specificity, with 83.3% sensitivity and 88.9% specificity in a large, 66-drug clinical dataset. The CNS-3D model was not directly tested on external datasets because either those studies did not collect time-series data in a format compatible with the feature-extraction workflow or the data were not available. However, the published comparator values reflected the best reported performance for each platform, and the CNS-3D model showed superior balanced sensitivity and specificity in this benchmarking analysis. Together, these findings indicate that functional profiling of CNS-3D organoids, when paired with clinically grounded drug selection and machine learning, provides improved prediction of seizure liability relative to existing *in vitro* preclinical approaches.

**Figure 5.**
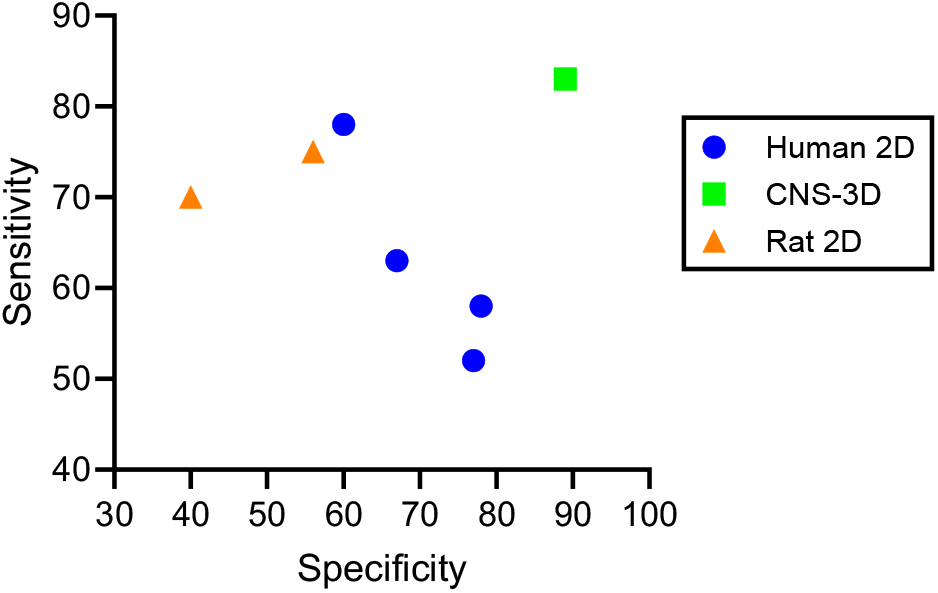
CNS-3D seizure-liability predictions benchmarked against published *in vitro* models. Sensitivity and specificity of the CNS-3D neurotoxicity prediction model compared with published seizure-liability models using MEA, calcium oscillation, and ion-channel assay platforms across human and animal-derived systems. Each point represents the performance of a reported model or platform. CNS-3D performance is shown at 83.3% sensitivity and 88.9% specificity.

## Discussion

Seizure liability remains a difficult safety endpoint to anticipate before human exposure, in part because seizures arise from perturbations in integrated neuronal network function rather than from a single molecular initiating event. In this study, we evaluated whether calcium activity in human iPSC-derived CNS-3D organoids could be used to predict clinically observed seizure risk across a retrospective, pharmacokinetically anchored drug set. The final model achieved high balanced performance, with 83.3% sensitivity and 88.9% specificity, supporting the use of CNS-3D functional profiling as a human-relevant component of neuro-safety assessment.

A central feature of the platform is that CNS-3D organoids are generated by iPSC co-differentiation rather than post hoc aggregation of terminally differentiated neural cell populations, an approach used in other HTS-compatible neural spheroid systems^12^. This distinction may be important for neurotoxicity applications because co-differentiation allows interacting neural populations to emerge within the same three-dimensional culture following normal developmental processes^19,20^. In the present study, CNS-3D organoids contained defined neuronal and glial populations, expressed neuroactive receptor and ion-channel gene programs that aligned closely with adult cortex, and generated coordinated calcium oscillations that could be measured in a 384-well, FLIPR-compatible format.

The responses to canonical convulsants illustrate why seizure prediction is unlikely to be captured by a single engineered metric such as increased peak frequency^7,21,22^. 4-aminopyridine, bicuculline, and kainic acid each altered calcium activity, but did so with distinct waveform-level effects. This observation motivated the use of more advanced time-series feature extraction and machine learning rather than a rules-based classification scheme^23^. The resulting model used dose-dependent calcium responses as the primary predictive signal, with chemical features providing additional molecular context. Although SMILES-derived features contributed only modestly to performance in the current dataset, they were retained because chemical structure creates a bridge to public toxicology and cheminformatics resources, including toxicophore-based approaches and molecular explanation frameworks^24^. This integration may become more valuable as larger annotated datasets become available.

The clinical anchoring of the drug set is also important. Many prior *in vitro* seizure-liability studies have relied heavily on tool compounds, mechanistic reference drugs, or limited negative-control sets. Here, drugs were selected based on documented human outcomes, defined chemical structures, pharmacokinetic information, and successful CNS-3D activity measurement. Testing across approximately 0.1x to 100x clinical C_max_ allowed activity changes to be interpreted relative to human exposure rather than only nominal *in vitro* concentration. Future incorporation of additional biological modules that capture blood– brain barrier transport, hepatic metabolism, protein binding, immune signaling and other systemic determinants of exposure could further improve predictive performance beyond the 83.3% sensitivity and 88.9% specificity reported here. Such integrated organoid and organ-on-chip systems are an active area of development, including multi-organ platforms linked by vascular flow and approaches that combine organoids with organ-on-chip engineering to improve drug-testing relevance^25^.

The comparison with published platforms suggests that human CNS-3D organoid calcium activity, when paired with clinically grounded machine learning, can improve balanced seizure-liability prediction relative to existing *in vitro* and preclinical approaches. This does not imply that CNS-3D functional profiling should replace all existing safety pharmacology assays. Rather, the results support its use as part of a weight-of-evidence framework, particularly where animal findings, ion-channel data, or 2D neural assays provide ambiguous or discordant signals. For example, one potential reason for 2D neuronal monocultures falling short is that they lack the multicellularity and network activity of the human brain. In contrast, CNS-3D organoids have multiple neuronal and glial cell types developing together that result in highly coordinated network activity. Thus, similar to seizure-related brain network disruption, perturbation of the network activity in CNS-3D is detected as a seizure risk. The ability to detect such risk with the high specificity observed here is especially relevant for decision-making, because false-positive seizure flags can halt otherwise promising programs or drive unnecessary follow-up studies.

Overall, these findings support CNS-3D organoid calcium activity as a scalable, human-relevant functional readout for seizure-liability prediction. Prospective analysis of new drugs will be valuable for defining real-world performance and could expand the current training set and enable iterative model improvements. When combined with clinically curated drug annotations, exposure-aware experimental design and machine learning, CNS-3D profiling provides a practical approach for identifying neurotoxicity risk earlier in drug development and for contributing functional human data to broader safety decision-making frameworks.

## Methods

### CNS-3D organoid generation and culture

CNS-3D organoids were generated from human iPSC-derived neural progenitor cells (NPCs), which were generated as previously described^9^. NPCs were maintained in Corning DMEM/Ham’s F-12 50/50 medium supplemented with B-27 Supplement (50x; Thermo Fisher Scientific, 17504044) and N-2 Supplement (100x; Thermo Fisher Scientific, 17502-048) at 37°C and 5% CO_2_.

For organoid generation, NPCs were seeded in 384-well, round-bottom, ultra-low attachment, spheroid microplates (Corning, 4516) at 30,000 cells per well. Forty-eight hours after seeding, organoids were maintained in BrainPhys Complete Medium consisting of BrainPhys Neuronal Medium and SM1 supplement (STEMCELL Technologies, 05792), supplemented with 20 ng/mL brain-derived neurotrophic factor (BDNF; STEMCELL Technologies, 78005), 20 ng/mL glial cell line-derived neurotrophic factor (GDNF; STEMCELL Technologies, 78058), and 1x HyClone Penicillin–Streptomycin (Cytiva, SV30010). Half-media changes were performed three times weekly at 48–72-hour intervals. Cultures were maintained at 37°C and 5% CO_2_.

After 6 weeks of differentiation, every batch of organoids underwent several quality control checks. These included mycoplasma testing, microbial sterility testing, and qualitative assessment of cellular composition by immunostaining for microtubule-associated protein 2 (MAP2; neurons), glial fibrillary acidic protein (GFAP; astrocytes), and neuroepithelial stem cell protein (Nestin; NPCs). Baseline calcium activity measurements over a 10-minute recording window were also included. Organoids used for drug screening were assayed during weeks 9 and 10 of differentiation.

### Immunocytochemistry

Organoids bound for immunostaining were transferred to high-content imaging PhenoPlate 384-well, black, optically clear, flat-bottom plates (Revvity, 6057302). Organoids were fixed overnight in PBS with calcium and magnesium (PBS+/+) containing 4% paraformaldehyde, permeabilized for at least 1 hour at room temperature in PBS+/+ containing 0.1% Triton X-100 (Sigma-Aldrich, X100-500ML), and blocked in PBS+/+ containing 0.1% Triton X-100 and 1% BSA Fraction V (Thermo Fisher Scientific, 15260037). Primary antibodies were diluted in blocking buffer and incubated overnight: guinea pig anti-MAP2 (Synaptic Systems, 188 004; 1:1,000), chicken anti-GFAP (Novus Biologicals, NBP1-05198; 1:1,000), and mouse anti-Nestin (BD Biosciences, 561230; 1:500). Secondary antibodies were diluted in blocking buffer and incubated overnight: Alexa Fluor 488 goat anti-guinea pig (Thermo Fisher Scientific, A-11073), Alexa Fluor 568 goat anti-mouse (Thermo Fisher Scientific, A-11031), and Alexa Fluor 647 goat anti-chicken (Thermo Fisher Scientific, A-21449). Nuclei were stained for 5 min with Hoechst 33342 (Thermo Fisher Scientific, H3570; 1:3,000).

### Fluorescence imaging

Fixed and stained organoids were imaged using a Yokogawa CQ1 high-content imaging system coupled with CQ1 Voyager software. Plates were first pre-scanned using a 4x dry objective to identify organoid positions. Identified regions were re-scanned using a 20x dry objective. High-magnification images were acquired as z-stacks with 10-µm intervals, 21 slices per field, and maximum-intensity projections were calculated for each channel. Images were acquired using 405-nm excitation with BP447/60-nm emission, 488-nm excitation with BP525/50-nm emission, 568-nm excitation with BP617/73-nm emission, and 640-nm excitation with BP685/40-nm emission.

### Bulk and single-nucleus RNA-seq

For bulk RNA-seq, mRNA was extracted using the Zymo Quick-RNA Microprep Kit (Zymo Research, R1050) from pooled material consisting of n=8 CNS-3D organoids or approximately 1×10^6^ NPCs per sample. RNA-seq data generation was performed by Novogene using Illumina 150-base pair (bp), paired-end sequencing. The analysis workflow included raw-read quality control using FASTQC, alignment to the human GRCh38 reference genome using *STAR*, expression quantification using *featureCounts*, and expression normalization using *EdgeR*. Reported expression values represent transcripts per million (TPM). CNS-3D organoids collected at 6, 8, and 10 weeks of differentiation were compared with source iPSCs, NPCs, adult human cortex mRNA (Takara, 636561), and whole fetal brain mRNA (Takara, 636526) by principal component analysis (PCA).

For single-nucleus RNA-seq, nuclei were isolated from flash-frozen CNS-3D organoids and 3′ gene expression libraries were generated using the 10x Genomics Chromium platform with Next GEM v3.1 chemistry. Libraries were sequenced on an Illumina NovaSeq platform with paired-end 150-bp reads, and clean reads were processed and normalized with Cell Ranger v6.1.2 using the human GRCh38 reference genome. Downstream analysis was performed in R using *Seurat*. Nuclei were filtered based on detected features and mitochondrial read percentage, mitochondrial genes were removed, and datasets were normalized using *SCTransform* with *glmGamPoi*. Integrated objects were generated using *Seurat* anchor-based integration, followed by PCA, Uniform Manifold Approximation and Projection (UMAP) dimensionality reduction visualization, nearest-neighbor graph construction, clustering, and marker-gene-based cell-type annotation.

### Calcium flux assay

Functional activity was measured using fluorescence-based calcium imaging on a FLIPR Tetra High Throughput Cellular Screening System (Molecular Devices). On the day of the assay, maintenance medium was exchanged for BrainPhys Phenol Red-Free Complete Medium consisting of BrainPhys without phenol red (STEMCELL Technologies, 05791), SM1 supplement (STEMCELL Technologies, 05711), BDNF (STEMCELL Technologies, 78005), GDNF (STEMCELL Technologies, 78058), and penicillin–streptomycin (Cytiva, SV30010). Three half-media exchanges were performed to remove phenol red, ending with 25 µL per well.

FLIPR Calcium 6 dye Component A (Molecular Devices, R8190) was equilibrated to room temperature and reconstituted in 11 mL of BrainPhys Phenol Red-Free Complete Medium. Reconstituted dye was added 1:1 to organoid wells by dispensing 25 µL per well, resulting in a 50 µL assay volume. Plates were incubated for 2 hours at 37 °C and 5% CO_2_ to allow for dye equilibration. For multi-plate experiments, dye addition was staggered across plates by 10 minutes.

The FLIPR instrument was equilibrated before acquisition with the stage set to 37°C. Calcium fluorescence was recorded using excitation/emission wavelengths of 470– 495/515–575 nm, excitation intensity of 40%, exposure time of 0.4 s, and camera gain of 40. Recordings were acquired every 0.5 seconds for 1,200 reads, corresponding to a 10-minute recording at 2 Hz.

Baseline spontaneous activity was recorded before drug addition. Drugs were then added as 10 µL of 6x dosing solution per well using the FLIPR liquid-handling system. After drug addition, plates were returned to 37 °C and 5% CO_2_ for at least 20 minutes between reads. Post-exposure recordings were acquired at 0.5, 1, 2, and 4 hours after drug addition; recordings beyond 4 hours were not used because prolonged Calcium 6 exposure may alter organoid function.

Each screening plate included vehicle controls, 30 µM 4-aminopyridine (Tocris, 0940) as an excitatory control, and 10 µM muscimol (Tocris, 0289) as an inhibitory control. Plates were required to show increased peak frequency after 4-aminopyridine treatment, suppression of spontaneous activity after muscimol treatment, and a Z′ factor ≥0.5 for vehicle versus muscimol control wells.

### Calcium trace processing and feature generation

Raw calcium traces were linked to plate, well, drug, concentration, replicate, and time-point metadata, then normalized to plate-matched vehicle controls. Feature extractors were used to quantify functional changes in calcium time-series recordings and drug molecular structures. Replicate traces were aggregated before generating drug-level predictions rather than generating independent replicate-level predictions and averaging model outputs. Concentration dependence was captured via dose-weighted trace averaging. Single-dose selection strategies were also evaluated during model development but were not used in the final workflow.

### Drug selection and clinical annotation

Drug selection and clinical annotation were performed as described in the study protocol^18^. Briefly, a retrospective, clinically-anchored drug set was assembled to evaluate prediction of human seizure liability. Drugs were eligible for inclusion if they had the following: 1) documented clinical outcomes related to seizure liability or absence of seizure liability, 2) a defined small-molecule structure represented by canonical SMILES, and 3) pharmacokinetic information (C_max_) sufficient for exposure anchoring.

Annotations were curated from human clinical sources, including regulatory documents, prescribing information, online clinical or pharmacology databases, clinical trial reports, and peer-reviewed academic literature. No formal hierarchy was imposed among these source types. Drug annotations were compiled by a clinical consultant and subsequently reviewed by two additional subject-matter experts. Labels were cross-checked against prior *in vitro* seizure-liability training datasets when applicable. Ambiguous, conflicting, or insufficiently supported seizure-liability reports were resolved conservatively by excluding the drug from the final training set. For each retained drug, the annotation dataset captured compound identity, clinical outcome label, C_max_, fraction unbound, seizure-liability context, seizure prevalence, SMILES code, and primary source reference (Supplemental Table 2). Seizure-liability context included whether the drug was associated with therapeutic-range seizure events, overdose-related seizures, or no documented clinical seizure liability. Seizure prevalence was categorized as none (safe), rare, moderate, or frequent as previously described^16^.

The final dataset contained 66 small-molecule drugs, including 30 seizure-associated drugs and 36 comparator drugs without documented clinical seizure liability (Supplemental Table 2). The underlying retrospective clinical dataset represented 120,551 patients. Each drug was tested across a seven-point, half-log concentration series spanning approximately 0.1x to 100x clinical C_max_.

### Model training and evaluation

XGBoost, a gradient-boosted decision tree classifier, was trained with full hyperparameter tuning to predict clinical seizure liability from integrated functional and chemical features. Model evaluation was performed using three-fold, drug-level cross-validation, with all wells, concentrations, and replicate measurements for a given drug confined to a single fold.

Out-of-fold predictions were generated for each drug and aggregated to a drug-level neurotoxicity score. Model performance was quantified by receiver operating characteristic (ROC) analysis. Area under the ROC curve (AUROC) was used as the primary performance metric. Sensitivity and specificity were calculated on out-of-fold drugs at the threshold selected by Youden’s J statistic. The 120-min post-exposure recording was selected for final analysis because it produced the strongest predictive performance among tested time points.

### Benchmarking against published seizure liability models

Published seizure liability models were identified from studies using MEA recordings, calcium oscillation assays, animal-derived neuronal cultures, human 2D neuronal cultures, ion-channel assays, and prior human 3D calcium-based systems^5-7,16,17^. Reported sensitivity and specificity values were extracted from each study and compared with the final CNS-3D model. Because comparator studies differed in drug sets, endpoint definitions, and threshold-selection strategies, comparisons were interpreted as field-level benchmarking.

### Statistical analysis

All statistical analyses were performed in GraphPad Prism 10.5.0. Unless otherwise indicated, analyses were performed at the drug level. Cross-validation splits were also performed at the drug level such that data from any single drug did not appear in both test and training sets. Calcium recordings from individual organoids were retained during feature generation and aggregated before drug-level prediction. ROC analysis, AUROC, sensitivity, specificity, and confusion matrices were calculated from out-of-fold predictions. Group comparisons were performed using one-way or two-way ANOVA, as appropriate, followed by Tukey’s *post hoc* test for multiple comparisons.

